# The eIF2α kinase Heme Regulated Inhibitor (HRI) protects the host from infection by regulating intracellular pathogen trafficking

**DOI:** 10.1101/142406

**Authors:** Wael Bahnan, Justin C. Boucher, Petoria Gayle, Niraj Shrestha, Mark Rosen, Bertal Aktas, Becky Adkins, Arba Ager, Wasif N. Khan, Kurt Schesser

## Abstract

Phosphorylation of eIF2α by its kinases is a stress response universally conserved among the eukaryota. Previously, we reported that the eIF2α kinases Heme Regulator Inhibitor (HRI) and Protein Kinase R (PKR) control distinct activities of diverse bacterial pathogens. Specifically for *Listeria monocytogenes*, it was shown that in HRI-deficient cells there was a reduction in the translocation of the pathogen to the cytosolic compartment as well as reduced loading of pathogen-derived antigens on MHC-1 complexes. Here we show that *Hri* -/- mice, as well as wild-type mice treated with a HRI inhibitor, are more susceptible to listeriosis. In the first few hours of *L. monocytogenes* infection, *Hri* -/- mice supported greater pathogen proliferation in the liver compared to that observed in *Hri* +/+ mice. This greater susceptibility of *Hri* -/- mice was not due to deficits in immune cell development as proportions and numbers of innate and adaptive cell compartments were largely normal and these mice could mount potent antibody responses to a model T cell-dependent antigen. Using *in vitro* cellular infection assays, we show that the rate of pathogen efflux from infected *Hri* -/- macrophages and fibroblasts is significantly higher than that observed in infected *Hri* +/+ cells. In contrast to the stark differences between *Hri* +/+ and *Hri* -/- cells in the infection dynamics of virulent *L. monocytogenes*, HRI was entirely dispensable for killing non-virulent strains of *L. monocytogenes*. These results suggest that in wild-type cells, HRI helps ensure the cellular confinement of virulent *L. monocytogenes* and loading of cytosolic-derived antigens on MHC-1 complexes that limit pathogen spreading and activating innate immune responses, respectively.

---

It has recently been recognized that the translation rate of a substantial fraction of eukaryotic mRNAs are potentially regulated by the levels of phosphorylated eukaryotic initiation factor *2α* (eIF2α-P)^1,2^. In humans and in mice the levels of eIF2α-P are controlled by elF2a kinases (GCN2, HRI, PERK, and PKR) that are in turn activated by various ligands or stimuli. The heme regulated inhibitor (HRI) was first isolated in reticulocytes where it regulates the translation of β-globin-encoding mRNAs during erythropoiesis and was later shown to play a protective role against oxidative stress during erythroid differentiation^4,5^. HRI also mediates protective responses to a variety of stresses in non-erythroid cells as well as in yeast^6-10^. Protein Kinase R (PKR) is activated by dsRNA and acts to inhibit translation of viral-derived mRNAs primarily through its eIF2α kinase activity^11^.

We showed using a yeast-based screen that HRI genetically interacts with two different bacterial virulence factors^12^. We went on to demonstrate that in mammalian cells eIF2α phosphorylation occurs following infection and that this activation is absolutely required for the infection-induced activation of NF-κB signaling and expression of pro-inflammatory cytokines^13^ similar to what others reported for LPS stimulation^14^. Beyond cytokine expression, both HRI and PKR regulate key events upon cellular infection by a variety of bacterial pathogens. Specifically, cells lacking PKR are more susceptible to invasion by the bacterial pathogens *Yersinia*, *Chlamydia*, and *Listeria* whereas in cells lacking HRI the functioning of the type III secretion system (*Yersinia*), intracellular growth (*Chlamydia*), and trafficking to the cytosol (*Listeria*) are all greatly reduced^15^. For the latter pathogen, cytosolic invasion is considered a critical component of its virulence strategy. Here we show the unanticipated consequences of HRI deficiency in mice following infection with *Listeria* and provide a cellular mechanism that may contribute to their heightened susceptibility.

## RESULTS

### HRI-deficient mice are susceptible to listeriosis

*Hri* +/+ and *Hri* -/- mice^16^ were infected intravenously (IV) with various doses of *L*. *monocytogenes*. At the lowest dose tested (3000 colony-forming units, CFU) neither strain of mice displayed any obvious external signs of disease (*not shown*). At the intermediate dose tested (15000 CFU), *Hri* +/+ mice presented by day 3 with mildly ruffled coats but were otherwise normal (with one exception) in their movements and were healthy looking by day 14 (**Fig. 1A**). In contrast, *Hri* -/- mice infected with a dose of 15000 CFU started to present with ruffled coats and retarded movements by day 2 and all succumbed between day 3 and 6 (**Fig. 1A**). At the highest dose tested (150000 CFU; approximate LD50) both strains of mice displayed comparable levels of disease symptoms by day 2; however, all of the *Hri* -/- mice succumbed whereas half of the *Hri* +/+ survived this dose (**Fig. 1B**). These data indicate that HRI plays a protective role in the early phases of *L*. *monocytogenes* infection.

**FIGURE 1.**
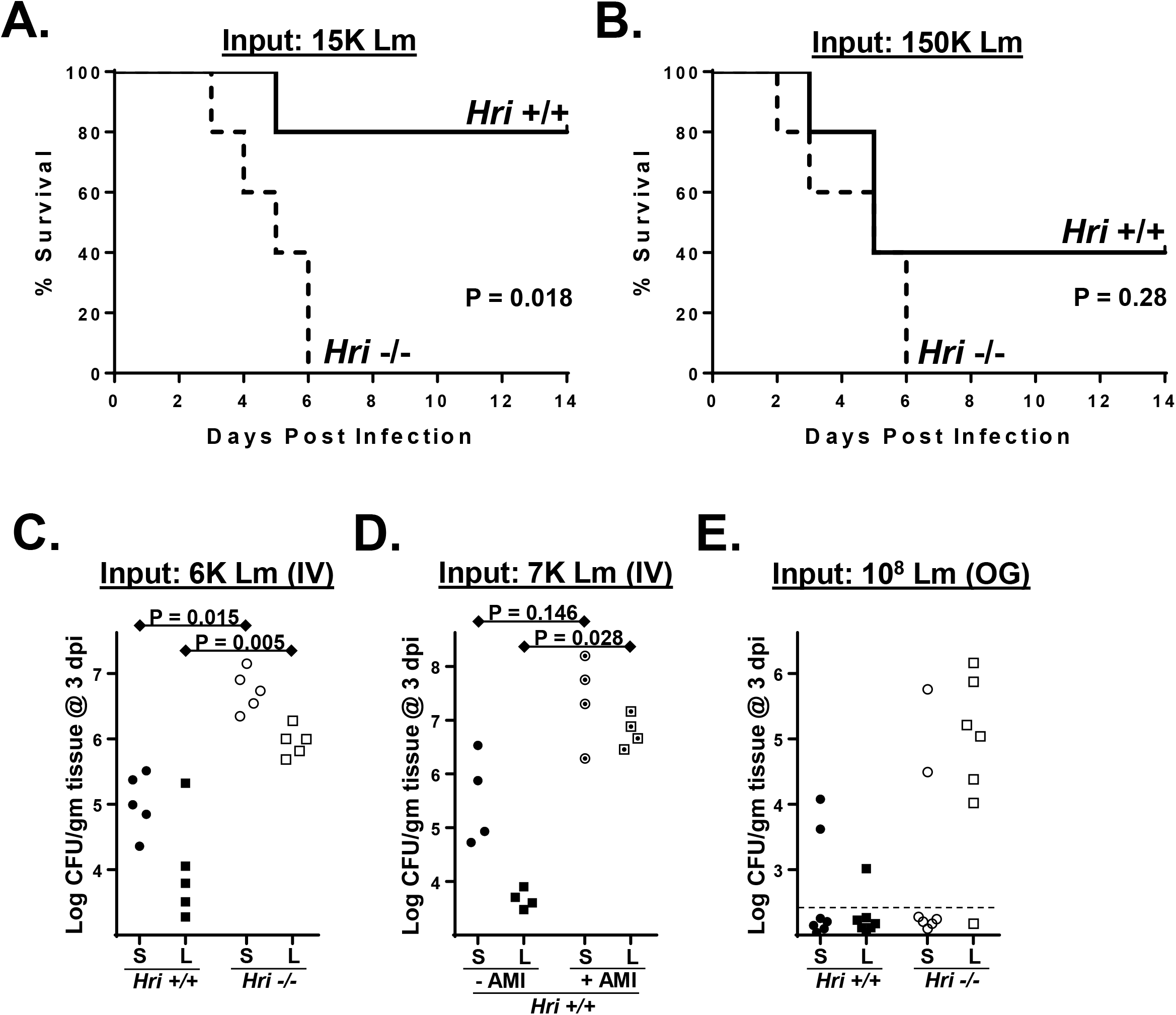
HRI is protective against listeriosis. *Hri* +/+ and *Hri* -/- mice (5 per group) were infected intravenously with 15000 **(A)** or 150000 **(B)** colony-forming units (CFU) of *L. monocytogenes* and monitored for 14 days. (P values by Mantel-Cox Log-rank test of a single representative experiment performed two times with similar results). **(C)** *Hri* +/+ (closed symbols) and *Hri* -/- mice (open symbols) were infected intravenously (IV) with *L. monocytogenes* and 3 days post infection (dpi) were humanely euthanized. Shown are the CFU in spleen (S, circles) and liver (L, squares) homogenates. **(D)** *Hri* +/+ mice infected with *L. monocytogenes* and treated with either vehicle (closed symbols) or the HRI inhibitor AMI (dotted open symbols) on days 0, 1, and 2 post infection. The load of *L. monocytogenes* in the spleen and liver was determined at 3 dpi. **(E)** Mice were inoculated orogastrically (OG) and at 3 dpi the load of *L. monocytogenes* in the spleen and liver was determined by CFU assay. Dotted line indicates the detection limit. (P values calculated using student *t* test of a single representative experiment performed two times.)

To determine whether increased disease susceptibility of HRI null mice was associated with enhanced pathogen colonization, bacterial loads in the spleen and liver were assessed in *Hri* +/+ and *Hri* -/- mice infected for 3 days with an intermediate dose of *L. monocytogenes*. There were significantly higher levels of viable *L. monocytogenes* recovered in the spleens (44-fold) and livers (21-fold) of *Hri* -/- mice compared to *Hri* +/+ mice (**Fig. 1C**). Recently a HRI inhibitor was identified in a high-throughput screen and was further modified to increase its specificity and bioavailability^17,18^. The resulting compound, here referred to as AMI (N-(2,6- dimethylbenzyl-6,7-dimethoxy-2H-[1]benzofuro[3,2-c]pyrazol-3-amine hydrochloride), disrupts HRI-dependent memory retention in the rat hippocampus as well as HRI-mediated activation of the β-site APP Cleaving Enzyme 1 (BACE1) in synaptic spines^19,20^. After 3 days infection with *L. monocytogenes*, among AMI-treated mice there was a greatly increased level of spleen (55-fold) and liver (1478-fold) colonization compared to untreated mice (**Fig. 1D**). These data show that a HRI null-like infection phenotype can be achieved pharmacologically in infected wild-type mice.

To further compare the innate responses of *Hri* +/+ and *Hri* -/- mice, an orogastric route of infection was used that mimics that of food-borne pathogens. Three days following being inoculated with *L. monocytogenes*, viable bacteria were recovered from the livers of only 1 of 7 *Hri* +/+ mice; in contrast, the livers of 6 of 7 *Hri* -/- mice had substantial bacterial loads following this infection period (**Fig. 1E**). Since the blood supply of the liver is primarily derived from the hepatic portal vein that drains the gastrointestinal tract, these data could indicate that HRI is important in front-line responses occurring in intestinal tissue. Collectively, these findings indicate that HRI plays a protective role against *L. monocytogenes* in a variety of infection scenarios.

### HRI restricts post-seeding pathogen proliferation in the liver

Relatively brief infection periods were used to distinguish between initial pathogen seeding of the liver by blood-borne *L. monocytogenes* and the subsequent intra-liver proliferation of the pathogen. Following IV infection of *L. monocytogenes*, the initial colonization of the spleen and liver occurred with comparable kinetics in *Hri* +/+ and *Hri* -/- mice. In both mouse strains ~10% of the inoculum is recovered in the spleen and liver 6 hours post infection (hpi) (Fig. 2A. However, by 24 hpi the recovery of *L. monocytogenes* from livers of *Hri* -/- mice were 10-fold higher than in *Hri* +/+ mice (Fig. 2A).

**FIGURE 2.**
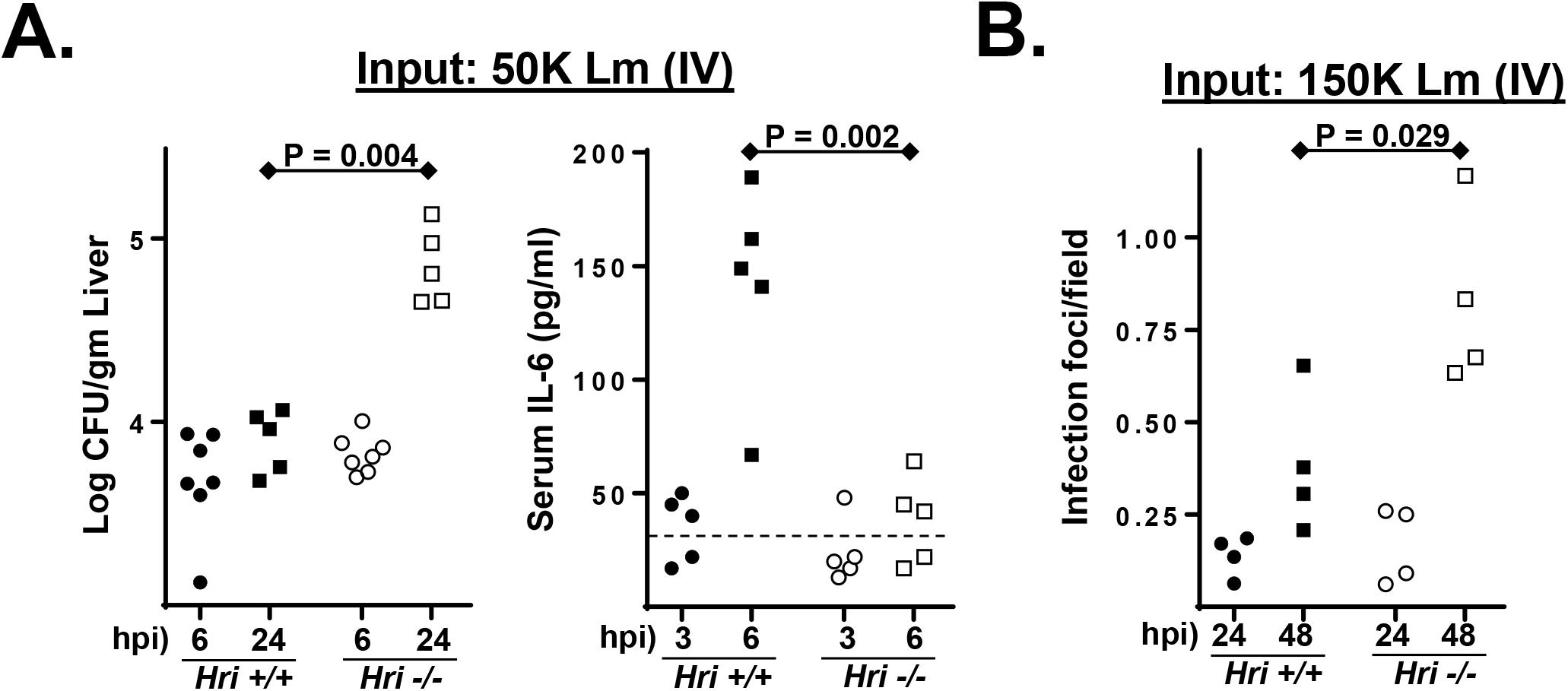
Enhanced post-seeding proliferation of *L. monocytogenes* in the liver is associated with delayed cytokine response. **(A)** *Hri* +/+ and *Hri* -/- mice were infected with *L. monocytogenes* and at either 6 or 24 hpi the bacterial load of the liver was determined by CFU assay. **(B)** At 6 hpi blood was collected and analyzed for IL-6 levels. Dotted line indicates the detection limit. **(C)** Histological analysis of livers from infected *Hri* +/+ and *Hri* -/- mice. Inflammatory micro-abscesses were enumerated by examining >20 microscopic fields for each of 4 individual mice per test condition. (P values calculated using student *t* test of a single representative experiment performed three times.)

Serum cytokine levels were measured in *Hri* +/+ and *Hri* -/- mice to determine whether immune activation differed in these strains. In both uninfected *Hri* +/+ and *Hri* - /- mice the levels of IL-6, TNFα, IFN-γ, MCP-1, IL-10, and IL-12p70 were below the level of detection. After 3 hours of infection, IL-6 was detected in the majority (3/5) of the *Hri* +/+ mice and by 6 hpi the levels had increased 3-fold (levels of the other cytokines remained undetected at 6 hpi) (**Fig. 2A**). In contrast, IL-6 was detected in only 1 of 5 *Hri* -/- mice after 3 hours of infection, and at 6 hpi IL-6 was minimally detected in 3 of 5 mice (**Fig. 2A**). Histological examination of livers from infected mice showed that at 24 hpi there were comparable levels of inflammatory foci in *Hri* +/+ and *Hri* -/- mice whereas by 48 hpi there were considerably greater number of these foci in the livers of *Hri* -/- mice (**Fig. 2B**). These data suggest that a delay in cytokine expression in *Hri* -/mice may contribute in allowing the pathogen to gain a foothold in the liver. The fact that inflammatory foci do form in these mice indicates that an appropriate response eventually occurs. These data are consistent with our previous findings showing defective cytokine expression in peritoneal and splenic *Hri* -/- macrophages infected *in vitro* with *L. monocytogenes*^15^. Collectively, these data suggest that events occurring subsequent to the initial pathogen seeding of these organs account for the differences observed at ≥ 24 hpi.

### HRI limits the emergence of virulent L. monocytogenes from macrophages

The *L. monocytogenes* infection cycle consists of four distinct steps: cellular invasion and initial enclosure within a *Listeria*-containing vacuole (LCV), virulence factor-mediated disintegration of the LCV membrane, replication within the cytosol, and finally the penetration of the plasma membrane resulting in infection of adjacent cells (as in an epithelial sheet) or, in solitary cells, release of the pathogen into the extracellular media that give rise to secondary infections. Peritoneal exudate macrophages (PEMs) derived from *Hri* +/+ and *Hri* -/- mice were used to determine whether HRI impacts the intracellular replication of *L. monocytogenes*. Freshly isolated *Hri* +/+ and *Hri* -/- PEMs were infected *in vitro* for 30 minutes, noninternalized bacteria were killed with gentamicin, and then viable intracellular bacteria were enumerated in cells infected for a total of 1 or 18 hours. Comparable levels of *L. monocytogenes* were recovered from *Hri* +/+ and *Hri* -/- PEMs following 1 hour of infection indicating that HRI does not play an apparent role in infectivity (**Fig. 3A**, *black bars*). Similarly, there were comparable recoveries of *L. monocytogenes* from *Hri* +/+ and *Hri* -/- PEMs following 18 hours of infection in the presence of gentamicin (**Fig. 3A**, *dark grey bars*). However, if gentamicin was removed from the infected cultures following the 30 minute killing phase, there was a 5-fold reduction in *L. monocytogenes* recovered from *Hri* +/+ PEMs compared to *Hri* -/- PEMs following 18 hours of infection (**Fig. 3A**, *light grey bars)* as well as a similar reduction in the bacterial levels in the overlying media (Fig**. 3A**, *unfilled bars*) suggesting that HRI impacts an extracellular phase of the *L. monocytogenes* infection cycle. A shorter-term kinetic analysis was performed to further characterize *L. monocytogenes* emergence from infected PEMs. The emergence of *L. monocytogenes* from infected *Hri* +/+ and *Hri* -/- PEMs occurred with similar kinetics (beginning >3 hpi), however, the magnitude of the increase was significantly greater in *Hri* -/- compared to *Hri* +/+ PEMs (**Fig. 3B**). These data further indicate that in the absence of HRI, the cellular efflux rate of *L. monocytogenes* is enhanced.

**FIGURE 3.**
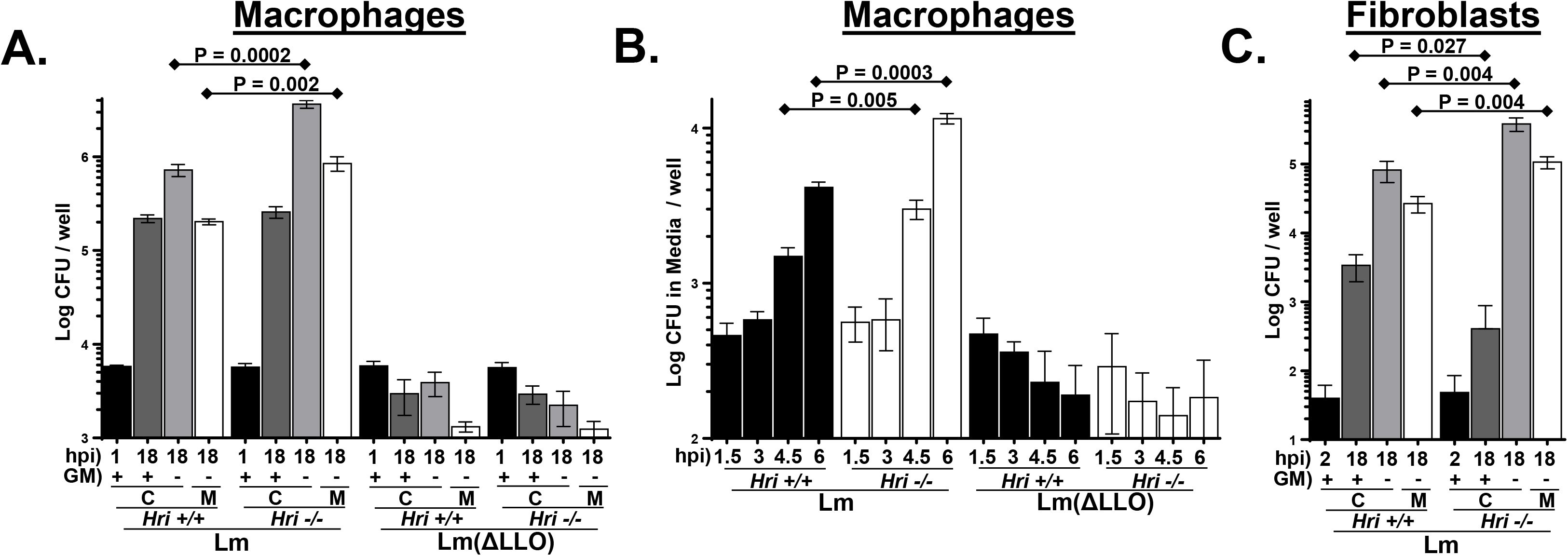
HRI restricts emergence of *L. monocytogenes* from infected macrophages. **(A)** Peritoneal exudate macrophages (PEMs) isolated from *Hri* +/+ and *Hri* -/- mice were infected *in vitro* with either wild-type *L. monocytogenes* (Lm) or a derivative mutant strain lacking the virulence factor listeriolysin O (ΔLLO). After 30 minutes of infection macrophages were treated with gentamicin for an additional 30 minutes to kill extracellular bacteria. Viable intracellular *L. monocytogenes* (C) were determined by CFU assay either at 1 hpi (black bars) or 18 hpi either in the presence or absence of gentamicin (dark grey and light gray bars, respectively). For 18-hr infections occurring in the absence of gentamicin (light gray bars), the levels of extracellular *L. monocytogenes* were determined in the overlying media (M, unfilled bars). **(B)** PEMs were infected as described above and following the removal of gentamicin at 1 hpi the levels of extracellular *L. monocytogenes* in the overlying media were determined at the indicated time points. **(C)** Mouse embryonic fibroblasts (MEFs) derived from from *Hri* +/+ and *Hri* -/- mice were infected with *L. monocytogenes* (Lm). After 90 minutes of infection cells were treated with gentamicin for an additional 30 minutes. Viable intracellular *L. monocytogenes* (C) were determined by CFU assay either at 2 hpi (black bars) or 18 hpi either in the presence or absence of gentamicin (dark grey and light gray bars, respectively). For 18-hr infections occurring in the absence of gentamicin (light gray bars), the levels of extracellular *L. monocytogenes* were determined in the overlying media (M, unfilled bars). (P values calculated using student *t* test of a single representative experiment performed > 3 times.)

*L. monocytogenes* lacking the virulence factor listeriolysin O (LLO), that is required for disintegration of the LCV membrane (discussed above), displayed a similar infection profile in *Hri* +/+ and *Hri* - /- PEMs. The infectivity (i.e., the initial invasion) of the *L. monocytogenes* (ΔLLO) strain was comparable to the LLO-expressing *L. monocytogenes* strain, however, in both *Hri* +/+ and *Hri* -/- PEMs the ΔLLO strain was subsequently efficiently killed and very few ΔLLO bacteria were detected in the overlaying media (**Fig. 3A** and **3B**). These data show that HRI is not required for the cellular defense against attenuated *L. monocytogenes*.

To determine whether HRI also acts to confine *L. monocytogenes* in fibroblastic cells, *Hri* +/+ and *Hri* -/- mouse embryonic fibroblasts (MEFs) were infected with *L. monocytogenes* and analyzed similarly as described above for macrophages. As was previously shown (ref. 15), *Hri* +/+ and *Hri* - /- MEFs are similarly infected by *L. monocytogenes* (**Fig. 3C**, *black bars*) and with prolonged infection in the presence of gentamicin, bacterial recovery from *Hri* +/+ MEFs actually exceeds (>8-fold) that of *Hri* -/- MEFs (**Fig. 3C**, *dark grey bars*). In this earlier study (ref. 15) the reduced bacterial recovery from *Hri* -/- MEFs was interpreted (as it turns out, erroneously) to be indicative that HRI was required for optimal *L. monocytogenes* intracellular proliferation. However, in the absence of gentamicin, bacterial recovery from *Hri* +/+ MEFs is reduced 5-fold compared to that of *Hri* -/- MEFs (**Fig. 3C**, *light grey bars*). Similar to macrophages, the increased levels of *L. monocytogenes* recovered from *Hri* -/- MEFs in antibiotic-free conditions are likely due to secondary infections since there was a 4-fold higher level of *L. monocytogenes* in the overlaying media of *Hri* -/- MEFs compared to *Hri* +/+ MEFs (**Fig. 3C**, *unfilled bars*). Collectively these findings show that in both macrophages and fibroblastic cells HRI promotes the cellular confinement of virulent *L. monocytogenes* that may account for the enhanced susceptibility of the *Hri* -/- mouse to this pathogen.

### HRI regulates intracellular trafficking dynamics

To determine whether HRI regulates intracellular trafficking in a non-infection setting, *Hri* +/+ and *Hri* -/macrophages were analyzed to determine whether they differed in their endosomal components or dynamics. PEMs derived from *Hri* +/+ and *Hri* -/- mice were phenotypically similar in terms of surface expression levels of macrophage-specific markers as well as the intracellular distribution of the early endosome marker EEA1 (**Fig. 4A**). In contrast, levels of the integral lysosomal factor LAMP-1 were notably higher in *Hri* -/- PEMs and were particularly concentrated in the perinuclear region (**Fig. 4A**). To compare dynamic responses, *Hri* +/+ and *Hri* -/- PEMs were pulsed with fluorescently-labeled beads. There were comparable levels of internalization of beads in *Hri* +/+ and *Hri* - /- PEMs, however, following their internalization, beads were much more rapidly trafficked to perinuclear regions in *Hri* -/- PEMs compared to *Hri* +/+ PEMs (**Fig. 4B**). These data reveal that HRI regulates the composition and dynamics of post-internalization trafficking and may account for the differential *L. monocytogenes* infection profile between *Hri* +/+ and *Hri* -/- macrophages.

**FIGURE 4.**
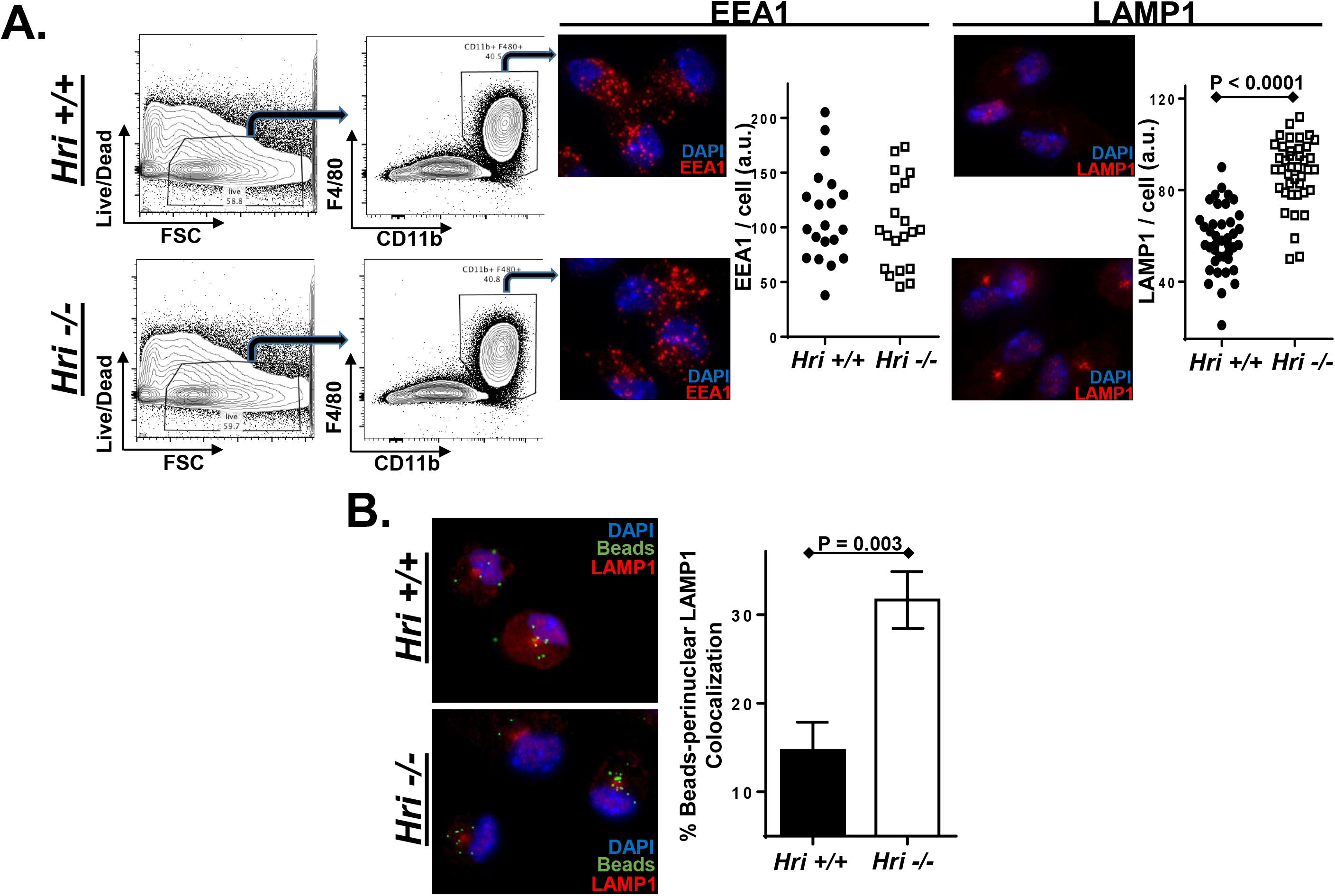
HRI regulates intracellular trafficking and lysosomal dynamics in macrophages. **(A)** PEMs isolated from *Hri* +/+ and *Hri* -/- mice were evaluated for viability (first column) and the gated population assessed for the macrophage markers F4/80+ and CD11b+ (second column). PEMs were stained for either EEA1 or LAMP1 and plotted is the total EEA 1- and LAMP 1- associated signal of individual cells. **(B)** *Hri* +/+ and *Hri* -/- PEMs were pulsed in vitro for 5 minutes with fluorescently-labeled beads and then analyzed 90 minutes later for bead localization and LAMP1 co-localization. Plotted is the percentage of beads localized to the LAMP1+ perinuclear region of three independent experiments in which >75 beads/experiment were tabulated. (P values calculated using student *t* test of a single representative experiment performed multiple times.)

### An eIF2 activator enhances HRI and PKR trafficking phenotypes

The above findings are drawn from HRI deficiency-based models, either genetic knockouts or inhibitors. To determine whether HRI regulated cellular activities could be hyperactivated, we employed a recently identified small molecule activator of eIF2α phosphorylation^21^. To determine whether this activator acted on post-internalization infection dynamics, macrophages were first infected and then treated with this eIF2α phosphorylation activator (I-17). This treatment regime resulted in a significant reduction in the intracellular proliferation of *L. monocytogenes* (**Fig. 5A**). If macrophages were first treated with I-17 and then infected, this resulted in a notable reduction in the infectivity (i.e., internalization) of *L. monocytogenes* (**Fig. 5B**). Macrophages pretreated with cycloheximide had comparable levels of infectivity as untreated cells (**Fig. 5B**), suggesting that the I-17 effect was not due to general translation inhibition. Additionally, I-17-treated macrophages were similarly resistant to internalizing inert beads as well as the soluble dye Lucifer yellow (**Fig. 5C**). Macrophages treated with I-17 had a highly punctate distribution of the early endosomal marker EEA1 (**Fig. 5D**) further indicating that I-17 disrupts early trafficking events. These findings are consistent with our previously published findings^13,15^ that HRI and PKR impact *L. monocytogenes* intracellular replication and pathogen invasion, respectively. Importantly, these data indicate that these separable cellular processes (invasion and post-invasion trafficking) can both be targeted by a small molecule activator.

**FIGURE 5.**
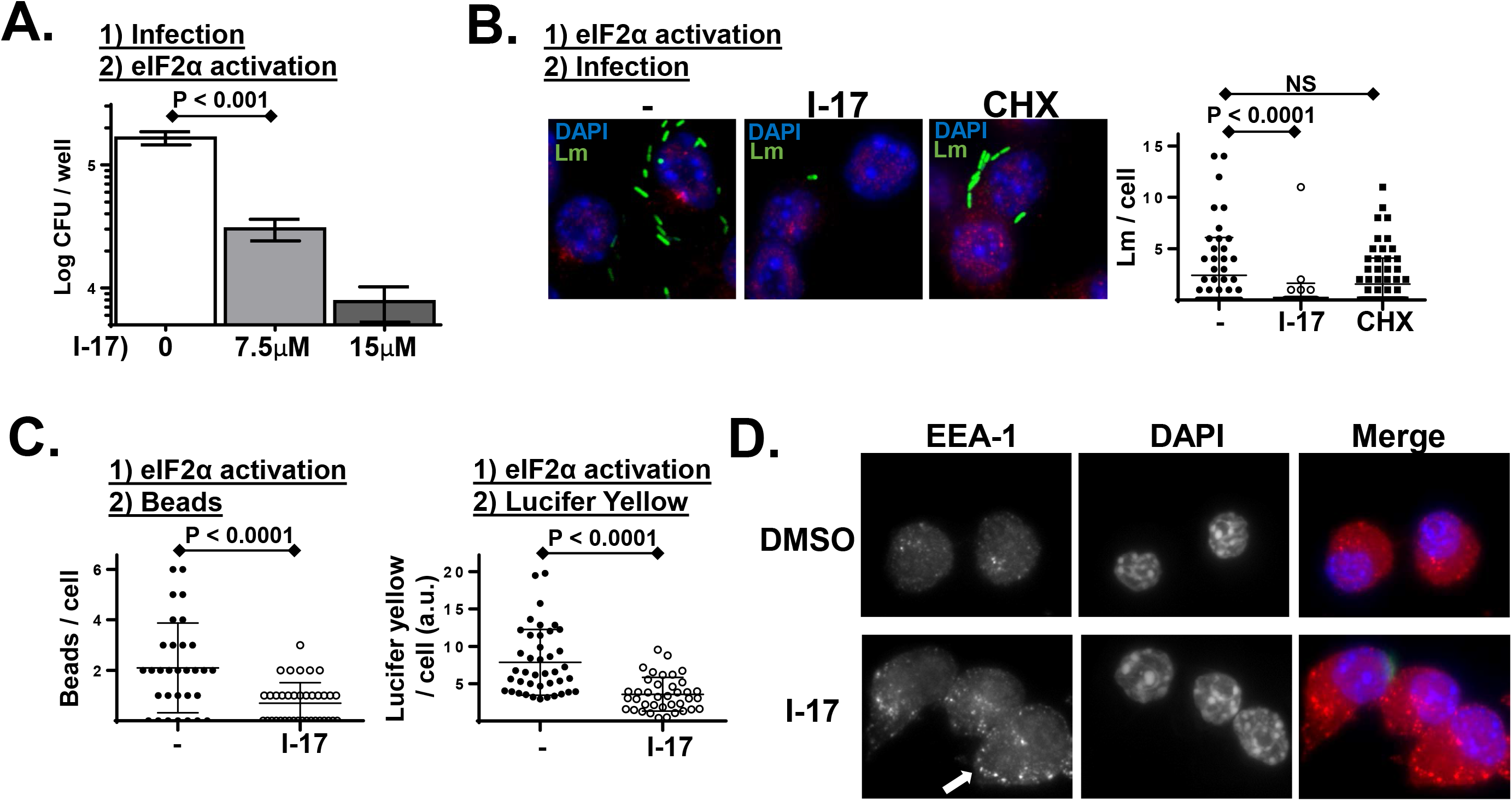
*L. monocytogenes* infection is inhibited in cells treated with a small molecule activator of eIF2α phosphorylation. **(A)** PEMs were infected with *L. monocytogenes* for 30 minutes, non-internalized bacteria were killed with gentamicin, and then cells were treated with the indicated concentration of the eIF2α phosphorylation activator I-17. Following 6 hours of infection, intracellular *L. monocytogenes* levels were determined by CFU assay. (P values calculated using student *t* test of a single representative experiment performed three times.) **(B)** Cultured murine macrophage-like cells (RAW 267.4) were either left untreated, treated with I-17,or treated with cycloheximide (CHX) for 60 minutes. Cells were then infected with GFP-expressing *L. monocytogenes* (Lm) for 45 minutes, washed extensively, and then analyzed by fluorescent microscopy. Plotted is the number of *L. monocytogenes* per cell from a representative experiment performed multiple times with similar results. **(C)** Similar experiment in which untreated and I-17-treated macrophages were pulsed with fluorescent beads (*left plot*)for 5 minutes or with the soluble dye Lucifer yellow (LY) for 20 minutes (*right plot*), washed extensively, and then analyzed by fluorescence microscopy. Plotted is the number of beads or fluorescence per cell from a representative experiment performed three times with similar results. **(D)** Similar experiment in which cells were treated with either vehicle alone or I-17 (10μM) for 60 minutes and then were fixed and stained for the early endosomal marker EEA1. The arrow indicates the cumulative early EEA1-positive endosomes in I-17-treated cells.

### Lymphoid and myeloid cell development is comparable between Hri +/+ and Hri -/- mice

The findings presented above and previously^15^ suggest that altered cellular pathogen trafficking, as well as delayed cytokine response may account for the enhanced susceptibility of HRI-deficient mice to *L. monocytogenes*. These early phases of immune responses against infection are regulated by innate immune cells. Therefore, additional investigations were performed to determine whether HRI impacted either the development of the immune system or the relatively longer-term adaptive immune responses. A comparative phenotypic analysis of immune cells isolated from the spleens of *Hri* +/+ and *Hri* -/- mice was performed by flow cytometry. There were no major differences between 21, 27, and 34 day old *Hri* +/+ and *Hri* -/- mice in their splenic cellularity or composition; both T and B lymphocytes and myeloid cells were comparable (data for 27-day old mice shown in **Fig. 6**).

**FIGURE 6.**
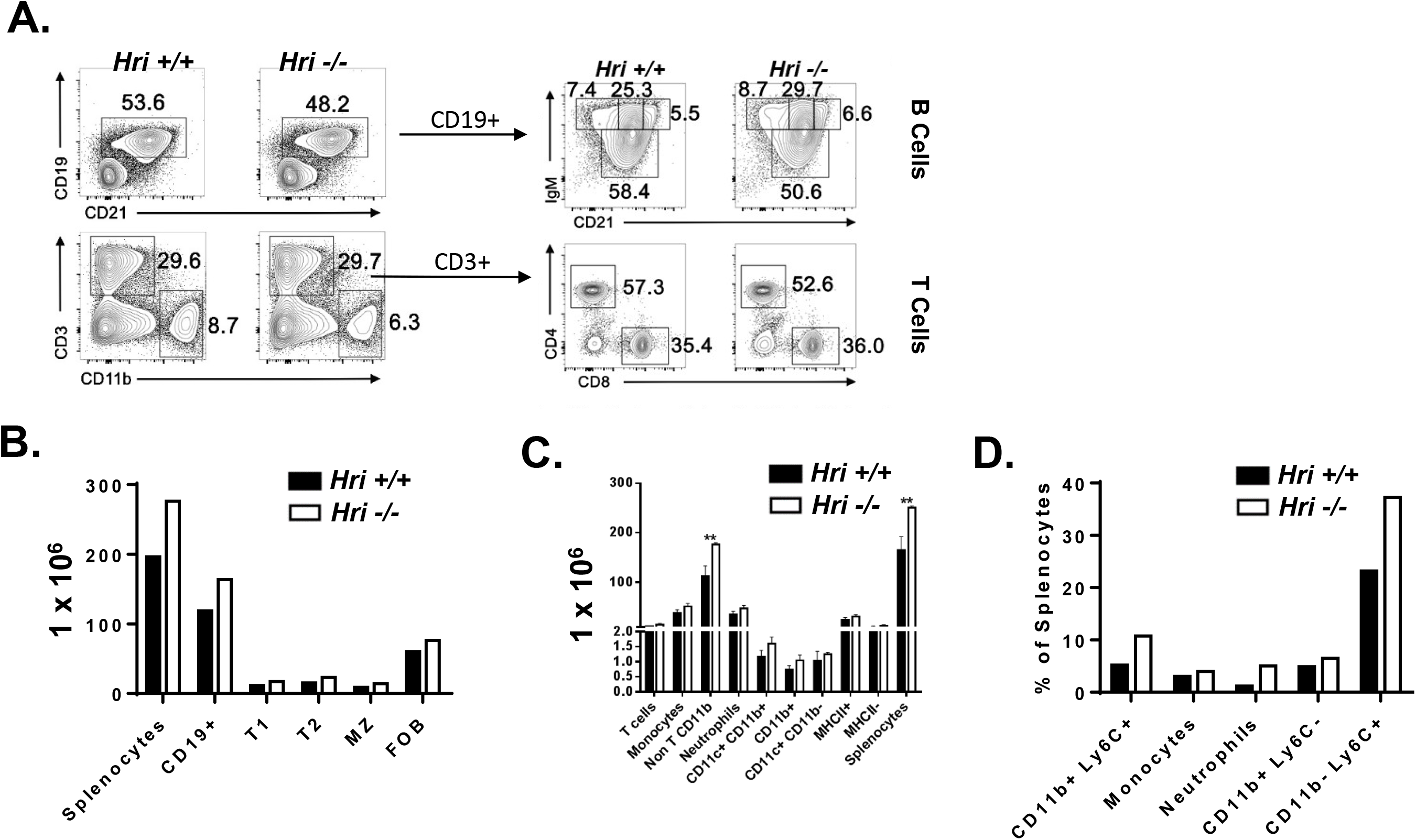
Largely intact immune cell development in HRI-deficient mice. Splenocytes isolated from 27-day old *Hri* +/+ and *Hri* -/- mice (N=4 each) were analyzed for lymphocyte and myeloid-derived cells by flow cytometry. **(A)** Plotted to the left is a representative mouse showing B, T, and macrophages and plotted to the right is higher resolution of B and T lineage subpopulations **(B, C)** the relative numbers of total splenocytes, total B cells and B cell subpopulations (B) and splenic T and innate cells resolved by separate cell surface markers **(C and D)** in an effort to reveal specific cell populations showing an effect of Hri deficiency. Shown is a representative experiment repeated multiple times with similar results.

To reveal any possible cell-autonomous effects of HRI deficiency on hematopoietic stem cell (HSC) development, we performed a bone marrow competitive chimera experiment in which 1:1 ratio of *Hri* +/+ and *Hri* -/- bone marrow cells (‘donors’) were transplanted into a lethally irradiated *Hri* +/+ recipient mice (**Fig. 7A**). After 5 weeks the reconstituted immune systems of the recipient mice were analyzed. There were no discernible differences in the ability of *Hri* +/+ and *Hri* -/- bone marrow-derived HSCs to contribute to the total number of B cells (B220+), B cell maturation (CD21), or CD3+ T cells or CD11b+ myeloid cells (**Fig. 7B**). Higher resolution analysis of B and T cells showed that all major subsets of splenic B cells including transitional (T1 and T2) and mature follicular and mature marginal zone B cells (FoB and MZ B) were present in both genotypes in similar proportions (**Fig. 7B**). Similarly, *Hri* +/+ and *Hri* -/- CD4+ and CD8+ T cells were in similar proportions (**Fig. 7B**). In addition to the ratios, the numbers of B, T, and myeloid cells were also comparable (**Fig. 7C**). These findings show that HRI does not play an apparent role in the development of immune cells and *Hri* -/- immune cells are not compromised in their ability to compete with *Hri* +/+ cells in the development of the immune system indicating that HRI is dispensable for the development of all major hematopoietic cell populations.

**FIGURE 7.**
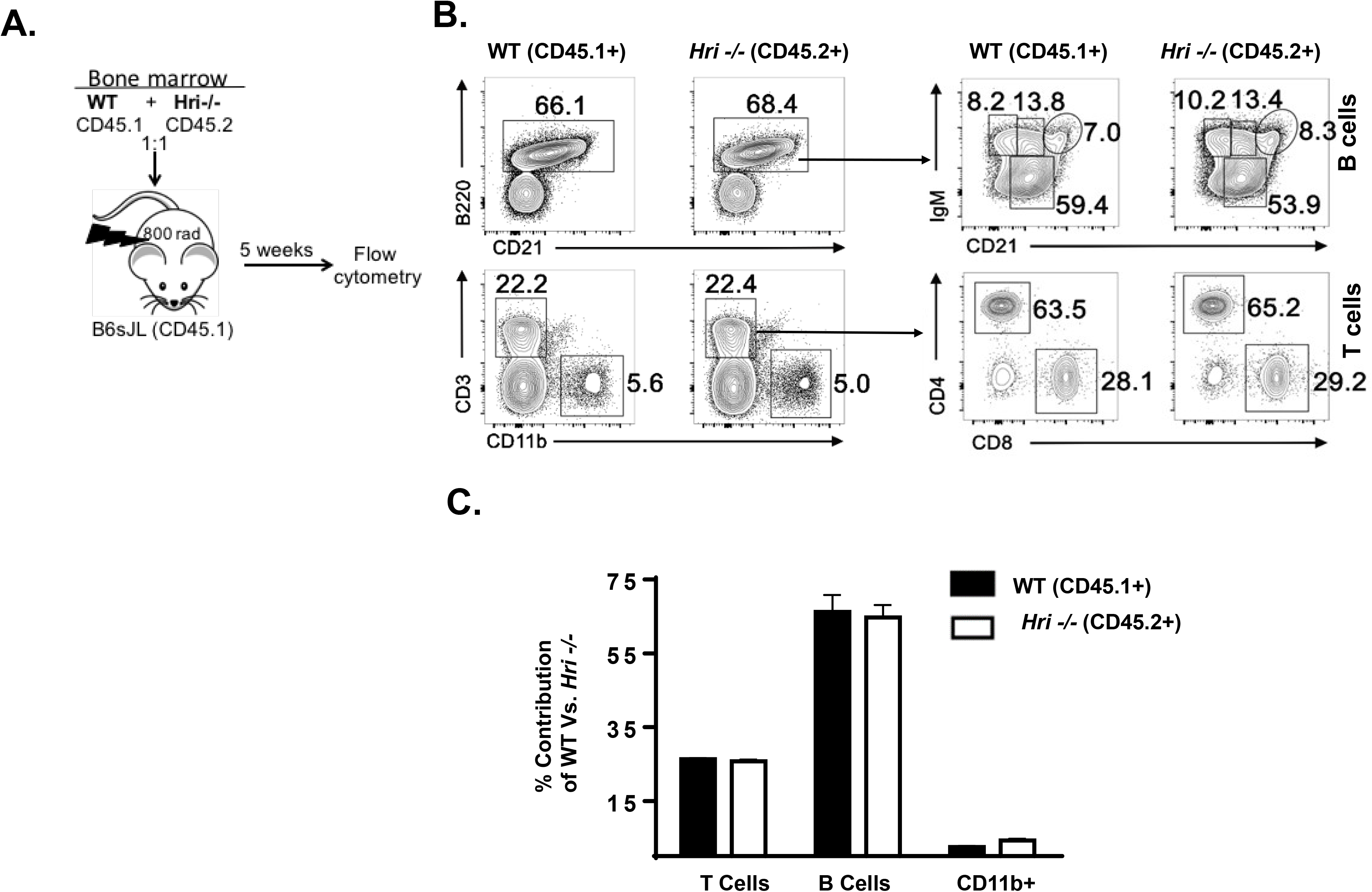
*Hri*-/- lymphoid and myeloid cells develop comparably in competition with *Hri* +/+ *in vivo.* **(A)** Bone marrow from wild-type and *Hri* -/- mice were co-transferred into congenic hosts. Splenocytes isolated from the recipient mice were analyzed after 5 weeks post-transfer to determine the percentages of the wild-type and *Hri* -/- derived B cells, T cells, and macrophages. **(B)** Plotted to the left is a representative mouse showing B, T, and macrophage populations and plotted to the right further resolved B and T cell subpopulations. **(C)** Bar graph representing percentages of T cells, B cells, and macrophages in recipient mice that were derived from wild-type or HRI deficient mice analyzed in (B). Shown is a representative experiment repeated twice with similar results.

### Increasedimmunoglobulin production and cellular expansion in Hri -/- mice

To determine any gross defects in the adaptive immune cell function of HRI deficiency, *Hri* +/+ and *Hri* -/- mice were tested for their capacity to mount an immune response to a T cell-dependent antigen (TNP-KLH). One week after immunization, *Hri* -/mice had detectably higher numbers of anti-KLH IgM antibody-secreting cells (ASCs) as well as in some mice increased IgG1 ASCs than that observed in *Hri* +/+ mice (**Fig. 8A, B**).

**FIGURE 8.**
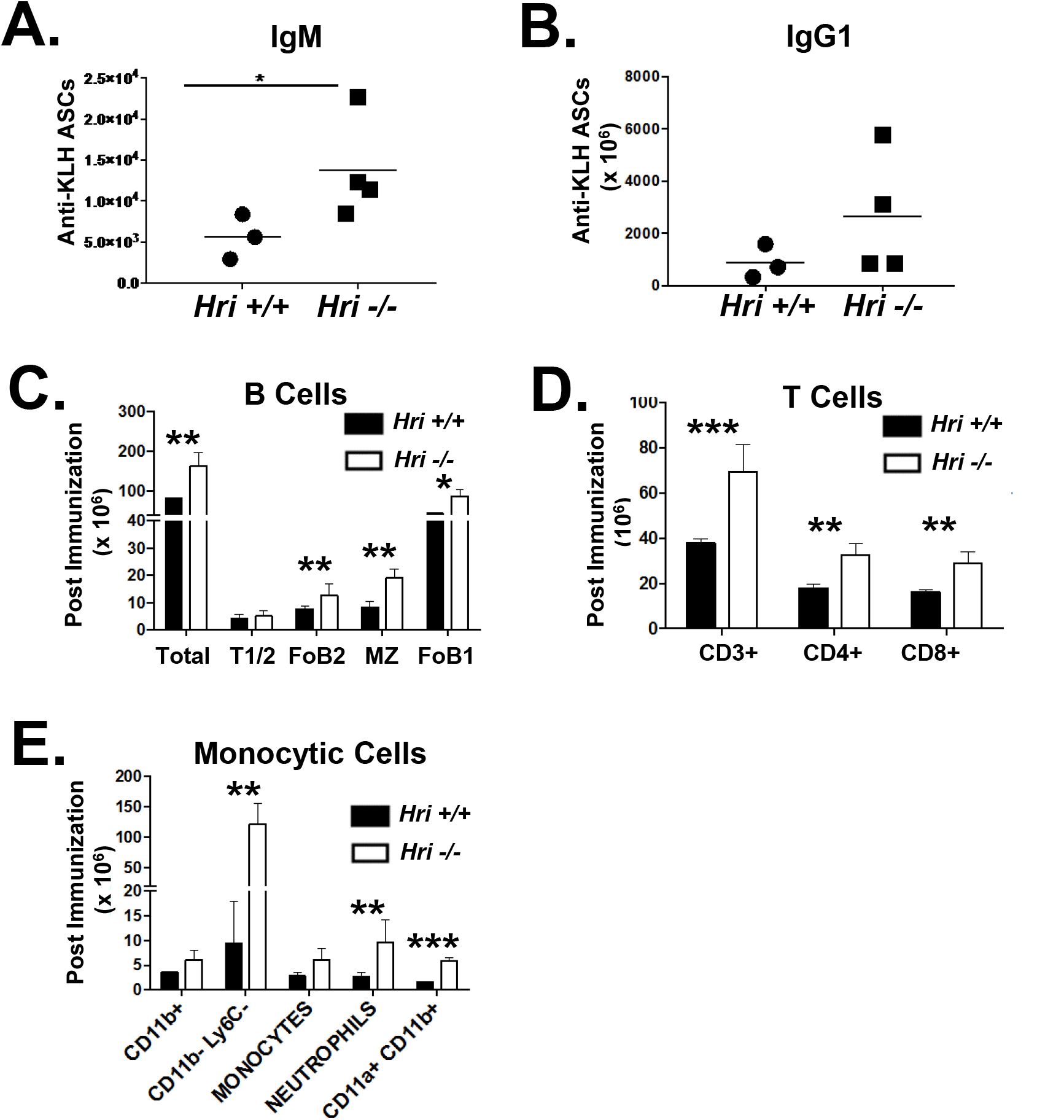
Largely intact adaptive immune cell development in HRI deficient mice following challenge with a T cell dependent antigen. **(A, B)** Frequency of IgM and IgG1 antibody secreting cells (ASCs) in response to the T cell dependent antigen TNP-KLH. Splenocytes from *Hri* +/+ and *Hri* -/- mice were analyzed by ELISpot for anti-KLH ASCs one week after challenge with TNP-KLH. The number of IgM **(A)** and IgG1 **(B)** ASCs are shown. Each data point represents an individual mouse. (C, D, E) One week after TNP-KLH immunization splenic B cell populations were enumerated for B cell subpopulations defined by CD19, IgM, IgD, AA4, CD21 and CD23 expression. **(C)** Splenic T and innate cell numbers were determined by FCM by staining for cell surface markers as indicated for T cells **(D)** and innate cells **(E).** Results shown from one experiment representative of three independent experiments with similar outcomes. *p< 0.05, **p< 0.01, ***p<< 0.001, ****p<0.0001.

The differences observed in antibody production led us to investigate cellular composition of the spleen. In *Hri* -/- mice the splenic B cell subsets (**Fig. 8C**), CD4+ and CD8+ T cells (**Fig. 8D**) as well as monocytic cells (**Fig. 8E**) were all increased in response to immunization compared to *Hri* +/+ mice indicating that while HRI is dispensable for the development of all major hematopoietic cell populations analyzed, it appears to restrain antigen-induced immune cell expansion and differentiation into ASCs. To further test whether HRI impacts adaptive responses, *Hri* +/+ and *Hri* -/- mice that had been infected with a sub-lethal dose (3000 CFU) of *L. monocytogenes* (see above) survived a subsequent challenge dose of 150 000 CFU administered 5 weeks later (*not shown*). These data indicate that HRI does not play an apparent role in generating adaptive responses.

## DISCUSSION

The enhanced proliferation of *L. monocytogenes* in *in vitro*-infected *Hri* -/- macrophages (~18 hpi) occurs with similar kinetics as the enhanced proliferation of this pathogen in the liver of *Hri* -/- mice. This concordance may indicate that the increased susceptibility of *Hri* -/- mice to *L. monocytogenes* infection is based on pathogen dynamics in initially-infected host cells. Our findings reported here and previously (ref. 15) indicate that HRI impacts the trafficking of *L. monocytogenes* in such a way that it limits the emergence or efflux of the pathogen from infected cells.

Screens to identify host genes that affect *L. monocytogenes* intracellular trafficking have found that factors involved in endosomal maturation can, when their expression levels are reduced, greatly impact infection^22-24^. Other studies have identified host factors that either neutralize the activity of Listerial-encoded virulence factors or can be exploited by the pathogen to promote infection. For example, the γ-interferon-inducible lysosomal thiol reductase (GILT) is required for the intracellular activation of listeriolysin O (LLO)^25^. Furthermore, we have recently shown that Perforin-2 inhibits the rapid acidification of *Listeria*-containing vacuoles preventing the virulence gene expression and activity^26^.

The findings presented here and earlier (in which it was directly shown that there is markedly reduced trafficking of *L. monocytogenes* to the cytosol in HRI deficient cells^15^) indicate that at the cellular level HRI and GILT are broadly similar in terms of their positive roles in promoting the vacuole to cytosol transitioning of *L. monocytogenes*. However, where these two host factors differ is in regard to the consequences that occur in their absence. The killing of *L. monocytogenes* is enhanced in GILT-deficient cells and GILT knockout mice are relatively protected against infection from this pathogen^25^. In contrast, here it is shown that the replication of *L. monocytogenes* is actually increased in HRI-deficient cells and that HRI knockout mice are relatively more susceptible to infection by this pathogen. The intracellular infection cycle in HRI-deficient cells that bypasses a cytosolic phase is characterized by a greatly enhanced rate of cellular efflux of the pathogen (**Fig. 3**). Reduced levels of cytosolic *L. monocytogenes* is usually correlated, as exemplified in the case of GILT, with reduced intracellular proliferation and disease. However, in the absence of a functioning immune system (i.e., the SCID mouse), vacuole-bound *L. monocytogenes* can cause chronic infections showing that at least under certain conditions cytosolic invasion is not invariably linked to disease^27^. Collectively our findings that modulation of intracellular pathogen trafficking pathways by the eIF2α kinases, HRI and PKR, suggests an important role for protein translational control in host response to infection and that these processes can be targeted with small molecule inhibitors and activators.

## EXPERIMENTAL PROCEDURES

*Mice and Mouse infections-Heme Regulated Inhibitor (Hri)* -/- mice (C57Bl/6), as well as the derivative *Hri* -/- mouse embryonic fibroblast (MEF) cell line, were generously provided by Jane-Jane Chen and Randal J. Kaufman, respectively^9,16^. The GFP-expressing wild-type *Listeria monocytogenes* 10403S strain and its ΔLLO derivative mutant strain were provided by Daniel Portnoy. Mice were treated humanely in accordance with all appropriate government guidelines for the Care and Use of Laboratory Animals of the National Institutes of Health and their use was approved for this entire study by the University of Miami institutional animal care and use committee (protocols 14-134 and 16-024). Seven-week old mice were infected intravenously with exponentially growing *L. monocytogenes* (O.D. 0.2) that had been propagated in brain heart infusion media at 37 °C. Infected mice were monitored for signs of distress daily and animals that presented reduced movements and/or ruffled coats were humanely euthanized. Livers were removed and homogenized in sterile water containing 0.05% triton X-100 by grinding through a fine wire mesh. Resulting homogenates were diluted further with water to fully lyse individual cells to release intracellular *L. monocytogenes* and the resulting dilutions plated to determine colony-forming units (CFU). Serum was collected as indicated in the figure legends and IL-6, TNFα, IFN-γ, MCP-1, IL-10, and IL-12p70 levels determined by a cytometric bead array assay (BD Biosciences). For developmental analyses, *Hri* +/+ and -/- mice were sacrificed on either day 21, 27, or 34. A single cell suspension of splenocytes were stained with CD19, CD11b, CD11c, CD4, CD8, CD3, and Live/Dead and analyzed by flow cytometry was using a LSRII cytometer (BD Biosciences). Chimera mice were generated using the recipient B6.SJL (congenic wild type) mice expressing the CD45.1 allele which were lethally irradiated with 1100 rads in a Gammacell 40 ^137^Cs irradiator (Best Theratronics). Sixteen hours later recipient mice were injected intravenously with 3.5x10^6^ bone marrow cells containing equal numbers of cells isolated from B6.SJL and *Hri -/-* donor mice. Six weeks after injection, mice were sacrificed and splenic leukocytes were analyzed as described above. Surface markers CD45.1 on B6.SJL and CD45.2 on *Hri* -/- cells were used to distinguish donor bone marrow cells.

*Cells and in vitro infections-* Peritoneal exudate macrophages (PEMs) were isolated from *Hri* +/+ and *Hri* -/- mice that had previously been injected with thioglycollate and assessed for viability using Live/Dead cell stain (Life Technologies) and analyzed as described^26^. Live cells were stained with antibodies APC clone M1/70 (Tonbo Bioscience) and PEcy7 clone BM8 (eBiosciences) to assess surface expression of macrophage-specific markers CD11b and F4/80. PEMs were seeded either at low density on glass coverslips (for microscopy) or into 48-well tissue culture dishes for cfu assays (2 x 10^5^ cells per well). In some experiments cells were treated with a freshly prepared solution of I-17 one hour after seeding. Two hours following their seeding, adherent PEMs were infected as described below. *Hri* +/+ and *Hri* -/- MEFs were seeded in 48-well tissue culture dishes (2 x 10^5^ cells per well) and infected the next day. Cultured cells were infected with exponentially growing *L. monocytogenes* (O.D. ~ 0.2) at a multiplicity of infection (MOI) of 5. At either 30 min (PEMs) or 60 mins (MEFs) post infection gentamicin (2 μg/ml; Gibco 15750-060) was added to kill extracellular *L. monocytogenes*. In control experiments it was determined that this concentration of gentamicin kills >99.992% of extracellular *L. monocytogenes* within 5 mins. Thirty minutes after the addition of gentamicin, cells were either lysed immediately (for early time points), left undisturbed (for later time points), or gentamicin-containing media removed and replaced with antibiotic-free media (to measure cellular *L. monocytogenes* efflux). Intracellular *L. monocytogenes* were determined by removing media from wells and adding 0.5 ml distilled sterile water for 30 sec, and plating the resulting water lysate (or dilutions thereof) on semi-solid LB media to enumerate CFUs following 24 hours incubation at 37 °C.

*Microscopy-* Following infections, macrophages bound to glass coverslips (see above) were fixed with 3.7% paraformaldehyde for 12 minutes and then stained for EEA1 (Abcam 2900) or LAMP1 (Abcam 24170) for 60 mins at room temperature. Macrophages were then washed and stained with AlexaFluor 555 for 60 mins prior to being washed and mounted onto precleaned glass slides. DAPI was included in the Pro-Long Gold mounting medium (Invitrogen). Imaging was performed using a Olympus fluorescence BX61 microscope was used that was equipped with Nomarski differential interference contrast (DIC) optics, a Uplan S Apo 100x objective (NA 1.4), a Roper CoolSnap HQ camera, a Sutter Lambda 10-2 excitation and emission filters, and a 175 watt Xenon remote source. Intelligent Imaging Innovations Slidebook 4.01 was used for image capture. A series of optical Z-sections (0.35 μm) were imaged and prior to analysis individual stacks were deconvolved using a nearest neighbor algorithm. Representative projected images were chosen to be included in the figures. ImageJ software was used to quantify the fluorescence signal per infected cell or per bacterium. The fluorescence signal was divided by the area of the cell or bacterium to generate a signal/area ratio that was termed ‘fluorescence intensity in arbitrary units’.

## Author contributions

WB and KS conceived the project. WB, JCB, and KS conducted the majority of the experiments, and together with WNK, analyzed and interpreted the results. PG and NS assisted in performing experiments. MR provided the HRI inhibitor and BA (Aktas) provided the elF2a kinase activator. BA (Adkins) and AA provided technical and material assistance. KS, with assistance from WNK, wrote the paper.

## FOOTNOTES

This work was supported in whole or part by National Institute of Health Grants R01 AI101041 and DK084246 (WNK) and R01 AI53459 (KS).

The abbreviations used are: HRI, heme regulated inhibitor; PKR, protein kinase R; PEMs, peritoneal exudate macrophages; hpi, hours post infection.

